# IMGT/RobustpMHC: Robust Training for class-I MHC Peptide Binding Prediction

**DOI:** 10.1101/2023.11.13.566840

**Authors:** Anjana Kushwaha, Patrice Duroux, Véronique Giudicelli, Konstantin Todorov, Sofia Kossida

## Abstract

The accurate prediction of peptide-MHC class I binding probabilities is a critical endeavor in immunoinformatics, with broad implications for vaccine development and immunotherapies. While recent deep neural network based approaches have showcased promise in peptide-MHC prediction, they have two shortcomings: (i) they rely on hand-crafted pseudo-sequence extraction, (ii) they do not generalise well to different datasets, which limits the practicality of these approaches. In this paper, we present PerceiverpMHC that is able to learn accurate representations on full-sequences by leveraging efficient transformer based architectures. Additionally, we propose IMGT/RobustpMHC that harnesses the potential of unlabeled data in improving the robustness of peptide-MHC binding predictions through a self-supervised learning strategy. We extensively evaluate RobustpMHC on 8 different datasets and showcase the improvements over the state-of-the-art approaches. Finally, we compile CrystalIMGT, a crystallography verified dataset that presents a challenge to existing approaches due to significantly different peptide-MHC distributions.

---

The critical molecular interactions between peptides and the Major histocompatibility complex (MHC) serve as the foundation for immune recognition which is the body’s ability to distinguish self from non-self and is pivotal in initiating immune responses against pathogens and malignant cells [60]. Accurate prediction of peptide-MHC (pMHC) binding is crucial in understanding the complexities of immune recognition and holds significant implications for the design of immunotherapies, vaccines, and personalized medicine [55, 61]. The MHC class I and class II proteins play a pivotal role in the adaptive branch of the immune system. T cells [11] play a vital role in the immune system by identifying specific peptides presented on cell surfaces through MHC molecules. This enables them to target and respond to threats, contributing to our body’s defense against infections and diseases. The MHC I molecules present peptides to CD8+ T cells, whereas the MHC II molecules present peptides to CD4+ T cells through the endogenous or direct and the exogenous or cross-presentation pathways [1, 9]. Specifically, the Human Leukocyte Antigen (HLA) system [18], a subset of MHC in humans, governs the presentation of peptides to T cells, thus initiating adaptive immune responses. HLA molecules are highly polymorphic, contributing to individualized immune responses and influencing susceptibility to various diseases [52]. Understanding HLA diversity and its implications is of paramount importance in fields ranging from immunology and transplantation medicine to vaccine development and disease association studies. This dynamic interplay between HLA molecules and immune responses underscores the central importance of HLA in human health.

Predicting peptide-MHC binding probability is a challenging computational task due to the vast combinatorial space of possible peptide sequences and the subtle nuances that govern their interactions with HLA molecules. Traditional approaches, such as scoring matrices and structural modeling, have made valuable contributions to this field. However, they often struggle to capture the intricate, high-dimensional relationships and the sequence patterns that are vital for accurate predictions. In recent years, machine learning, especially deep learning techniques have emerged as powerful tools for addressing this challenge. These methods leverage vast datasets of experimentally measured peptide-HLA binding probabilities to learn complex patterns and relationships. Nonetheless, existing models still face limitations in effectively encoding and representing the structural and sequential features of both peptide and MHC sequences.

In conventional approaches, predicting peptide-HLA binding probabilities has predominantly relied on two methods: scoring matrices [35] and structure-based modeling [38]. Scoring matrices assign scores to pairs of amino acids based on their co-occurrence frequencies in binding or non-binding interactions. Position-Specific Scoring Matrices (PSSMs) [30] refine this approach by considering the position of amino acids within a peptide. While these methods are computationally efficient, they may struggle with capturing subtle, high-dimensional relationships [39]. On the other hand structure-based modeling [45], leverages experimental three-dimensional structures of peptide-HLA complexes to analyze the physical interactions. This approach offers high accuracy, when structural data are available, but is limited by the availability of experimental structures[20, 29]. Both methods have played valuable roles in this field, but they may not generalize well to diverse peptide-HLA combinations or fully capture the complexity of binding probabilities [34].

Diverging from conventional approaches that often focus on peptide sequences, NetMHCPan [44] proposed a pseudo sequence encoding method to characterize the interactions between HLA and peptides, in which an HLA sequence is reduced to a pseudo amino acid sequence of length 34. Only 34 corresponding residues are encoded as input for an HLA sequence. This pseudo sequence representation-encoding model has been adopted by the rest of the other machine learning approaches except Deep-AttentionPan [19] which employs full HLA sequences, wherein each HLA sample is represented as 280 to 287 length residue sequence (8 to 15 from the peptide and 272 from the HLA). However, their performance is significantly worse as compared to pseudo-sequences based approaches like NetMHCPan. In NetMHCPan and other approaches a sample is represented as a 42 to 49-length residue sequence (8 to 15 from the peptide and 34 from the HLA).

In this research, we intend to harness the power of recent deep learning techniques, particularly efficient transformer models like Perceiver IO model [17], to predict peptide-MHC interactions [57]. To this end, we present PerceiverpMHC that leverages latent cross-attention design from Perceiver IO and therefore can scale up efficiently to longer sequences without requiring domain-specific architectural adaptations. This adaptability makes the PerceiverpMHC an ideal candidate for modeling the peptide-HLA binding phenomenon, as it can simultaneously consider the sequence information of both peptides and HLA molecules while capturing complex inter-dependencies. In our study, instead of focusing solely on peptide sequences, we take a broader approach by considering the full Major Histocompatibility Complex (MHC) sequences. By encompassing the entirety of the MHC molecule, including both the peptide-binding groove and the surrounding regions, our model gains a nuanced understanding of the structural context in which peptides bind. This holistic approach not only accounts for polymorphism within the MHC gene but also captures the dynamic interplay between residues that may influence peptide presentation on the surface of MHC molecule. therefore, our model is poised to provide more accurate and contextually relevant predictions of peptide-HLA binding probabilities.

While recent advances in deep neural networks have shown promise in peptide-HLA binding predictions, their performance tends to drop when confronted with datasets having slightly different distribution of peptide lengths which limits the applicability of these approaches. To address this issue, we enhance the robustness of PerceiverpMHC by incorporating the power of self-supervised learning, which we refer to as RobustpMHC. In contrast to traditional supervised learning relying solely on labeled data, self-supervised learning leverages both labeled and unlabeled samples to refine the model’s understanding of the underlying relationships. In the context of peptide-MHC binding, training RobustpMHC using self-supervision proves invaluable, as it allows us to harness a broader spectrum of data, augmenting experimentally validated interactions with synthetically generated data. Data augmentation not only enhances the robustness of our predictions but also enables the model to generalize more effectively to novel peptides and MHC sequences.

## Contributions

Previous architectures for peptide-MHC predictions rely on handcrafted pseudo-sequences, whereas natural language processing literature has proved that transformer architectures [23] can learn from long sequences. Therefore, we present a Perceiver IO based architecture (PerceiverpMHC) for predicting pMHC bindings on full HLA sequences. In contrast to existing approaches [2, 10, 31] that require hand-designed pseudo-sequences, we show that attention mechanisms can learn efficient representations enabling us to work with full sequences. Secondly, we observe that the performance of existing approaches significantly drops on datasets with slightly different distribution of peptide lengths. To mitigate this issue, we propose RobustpMHC that leverages self-supervised learning strategy resulting in superior performance over the state-of-the-art approaches across six different datasets. Thirdly, we introduce a new dataset CrystalIMGT dataset, a set of mass spectrometry observed peptide-MHC pairs, which presents a challenge to the existing approaches due to significantly different data distribution. Additionally, we propose a transfer learning technique that can substantially enhance performance using only 10% of the available data for the CrystalIMGT dataset. Furthermore, we demonstrate the capability of the features acquired through our method to transfer to the prediction of immunogenicity data for neoantigens. This transfer is assessed through a comparison with the nine state-of-the-art approaches. In conclusion, we present a robust pMHC binding prediction strategy and an exhaustive evaluation benchmark, which we believe will advance the field of pMHC binding prediction. Once this paper is accepted, we will make our implementation, trained models, and datasets available, to facilitate reproducibility. Additionally, we have deployed a user-friendly web interface for predicting peptide-MHC binding.

## 1 Results

We designed our experiments to investigate the following research questions:

1. Do we need to depend on hand-crafted pseudo-sequence extraction, or can neural networks autonomously discern crucial features?
2. How does PerceiverpMHC compare with other Transformer architectures, and how does RobustpMHC’s learning strategy compare against other self-supervised learning approaches such as data augmentation and contrastive learning?
3. To what extent do the introduction of mutations contribute to enhancing the robustness of RobustpMHC? How many mutations are sufficient, or does a large number of mutations degrade the performance?
4. Does RobustpMHC generalize well across diverse datasets, enhancing its practical applicability while effectively accommodating diverse peptide lengths in sequences?

To answer (1), we compare PerceiverpMHC with recent pseudo-sequence approaches and DeepAttentionPan (the single full sequence base approach) on 6 datasets in section 1.1.In section 1.3 we show that the learned representations of RobustpMHC can be transferred to datasets that are even significantly different than the training dataset. Finally, we present ablation studies in section 1.4 to answer the research questions (2-4). With regard to (4), we show that across 8 different datasets RobustpMHC is either state-of-the-art approach or at par with state-of-the-art approach for that dataset in section 1.2.

### 1.1 Full sequence evaluation

We initially evaluate whether neural networks can inherently learn which amino acids are crucial for binding given the full MHC sequence or if we need to design hand-crated pseudo-sequences. To this end, we evaluate the performance of PerceiverpMHC on four publicly available benchmarks: independent and external set from Anthem [33] dataset, Neoantigen [10] and HPV [8] datasets and compare with the state-of-the-art pseudo-sequences based approaches like TransPHLA [10], capsNet [21], NetMHCpan 4.1 [44] and BigMHC [2].

The independent set from Anthem dataset [33] contains 112 types of HLA alleles, whereas the external set contains five HLA alleles. The Neoantigen data [10] comprises non-small-cell lung cancer, melanoma, ovarian cancer and pancreatic cancer from recent works including 221 experimentally verified pHLA binders. The HPV dataset [10] 278 experimentally verified pHLA binders from HPV16 proteins E6 and E7, consisting of 8–11-mer peptides. Following TransPHLA [10] we report four metrics Area Under Curve (AUC), Accuracy, Matthews correlation coefficient (MCC) and F1 scores on independent set and external set. For the datasets such as Neoantigen and HPV that have only experimentally verified binders, we report the true positives and the false negatives during prediction. Furthermore, we compare our PerceiverpMHC approach with deep-AttentionPan [19], which is also a full sequence based deep pHLA binding prediction approach.

It can be seen from Figure 1a-c that even on full sequences PerceiverpMHC is at par with the state-of-the-art pseudo-sequence based approaches on three evaluation datasets. While PerceiverpMHC is slightly inferior on the HPV dataset (refer Figure 1d) as compared to TransPHLA and CapsNet, it is superior to other approaches including NetMHCPan4.1 and BigMHC-EL. This showcases that deep neural networks can implicitly learn to attend the important amino acids without external human supervision i.e. hand-crafted pseudo-sequences. Furthermore, in comparison to the previous full sequence method DeepAttentionPan [19], PerceiverpMHC achieves far better prediction performance, which clearly demonstrates the superiority of transformer based architectures. It is worth mentioning that TransPHLA has shown superior performance over 14 popular approaches therefore for brevity we do not show comparison with them [16, 19, 22, 25, 27, 31, 33, 40, 42, 50, 59, 62].

**Fig. 1.**
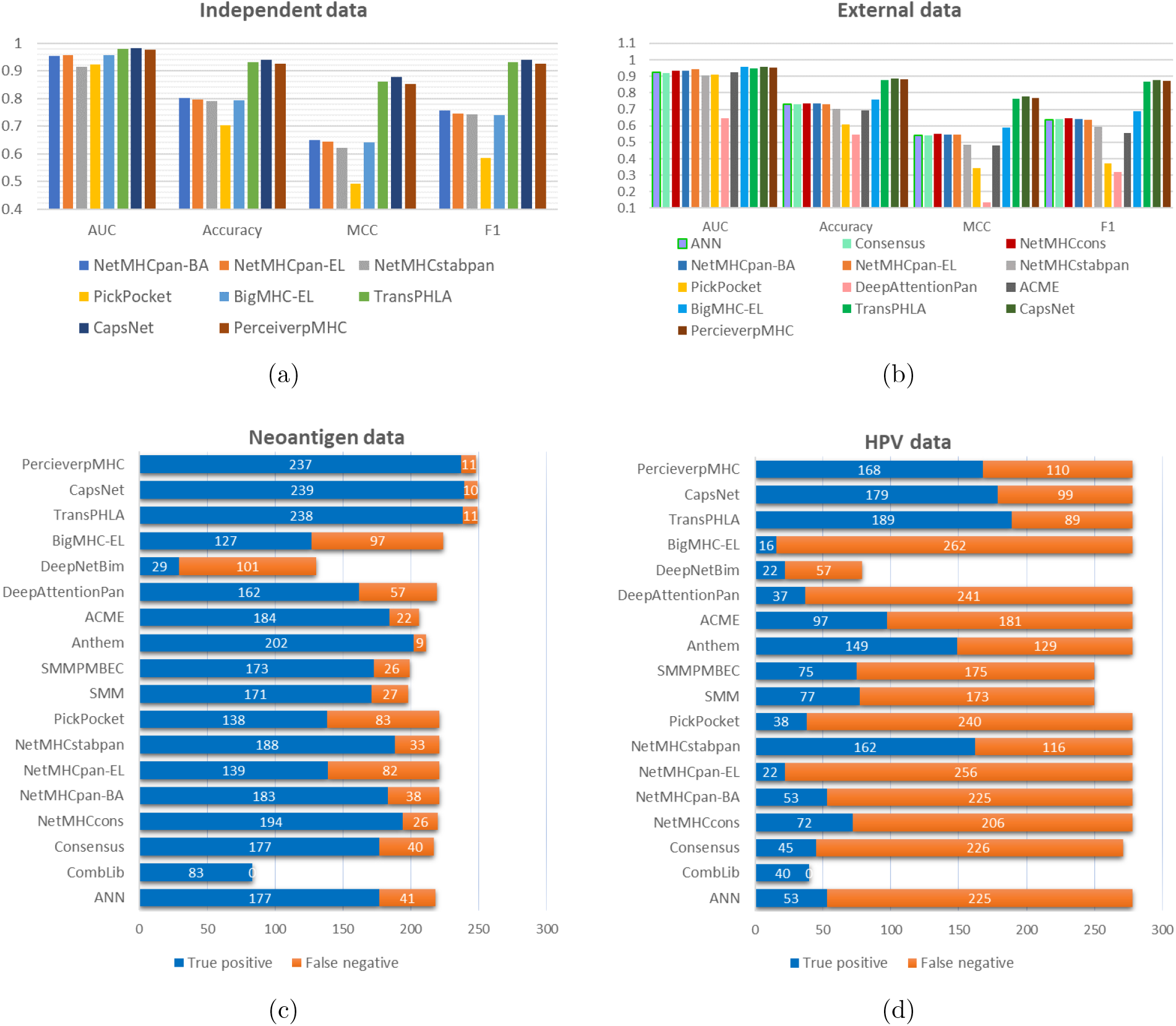
Full sequence evaluation. Comparison of PerceiverpMHC trained on full sequence with recent approaches on (a) Anthem Independent set, (b) Anthem External set, (c) Neoantigen data and (d) HPV data.

### 1.2 Robustness evaluation

We next evaluate how well our self-supervised robust training generalises across different datasets for peptide-HLA-I binding prediction. To this end, we evaluate RobustpMHC on six datasets, independent and external splits form Anthem datasets, HPV and Neoantigen dataset. We further compare them on Covid dataset and IEDB binary evaluation set. The current coronavirus disease 2019 (COVID-19) outbreak is a worldwide emergency, as its rapid spread and high mortality rate has caused severe disruptions. Understanding the immune patterns of COVID-19 is very crucial for mechanisms of SARS-CoV-2-induced immune changes, their effect on disease outcomes, and their implications for potential COVID-19 treatments [51]. The COVID dataset consists of 29 experimentally verified positive and 89 negative samples from different sources [32] [46].

The IEDB dataset [12] contains three different measurements IC50, T1/2 and binary. Here we consider the binary set which consists of 3845 experimentally verified EL(Mass spectrometry data include naturally presented MHC ligands, referred to as eluted ligands) [5] binding samples and 3656 negative samples. Similar to Anthem, IEDB has been one of the prominent publicly available benchmarks which contains slightly different distribution as compared to Anthem therefore it provides a diverse set for evaluating the generalization of the approaches.

It can be seen from Figure 2 that for independent set (Figure 2a), which has very similar peptide length distribution as training data, the performance of RobustpHMC is slightly lower as compared to CapsNet. In contrast, on external dataset (Figure 2b) RobustpHMC shows superior performance. Even on Neoantigen data RobustpHMC shows slightly better performance as shown in Figure 2c. On HPV dataset (Figure 2d) RobustpHLA is slightly behind TransPHLA but significantly better than CapsNet and NetMHCpan-4.1. On COVID dataset (Figure 2e) RobustpHLA is again superior, marginally outperforming NetMHCpan-4.1, and significantly outperforming TranspHLA and CapsNet. While on IEDB dataset (Figure 2f), RobustpHMC performs marginally better than CapsNet and TranspHLA and notably better than NetMHCpan-4.1. In conclusion, RobustpMHC obtains state-of-the-art results on four out of six datasets. Furthermore, RobustpHMC demonstrates consistent performance across all six dataset which showcases the robustness of our approach. It is worth noting that RobstpMHC and PerceiverpMHC work on full sequences while others baselines in Figure 2 require psuedo-sequences. Finally, RobstpMHC constantly out-performs PerceiverpMHC in all six datasets, therefore we can conclude that mutation robust training helps in generalization and can significantly improves the practicality of pHLA prediction approaches.

**Fig. 2.**
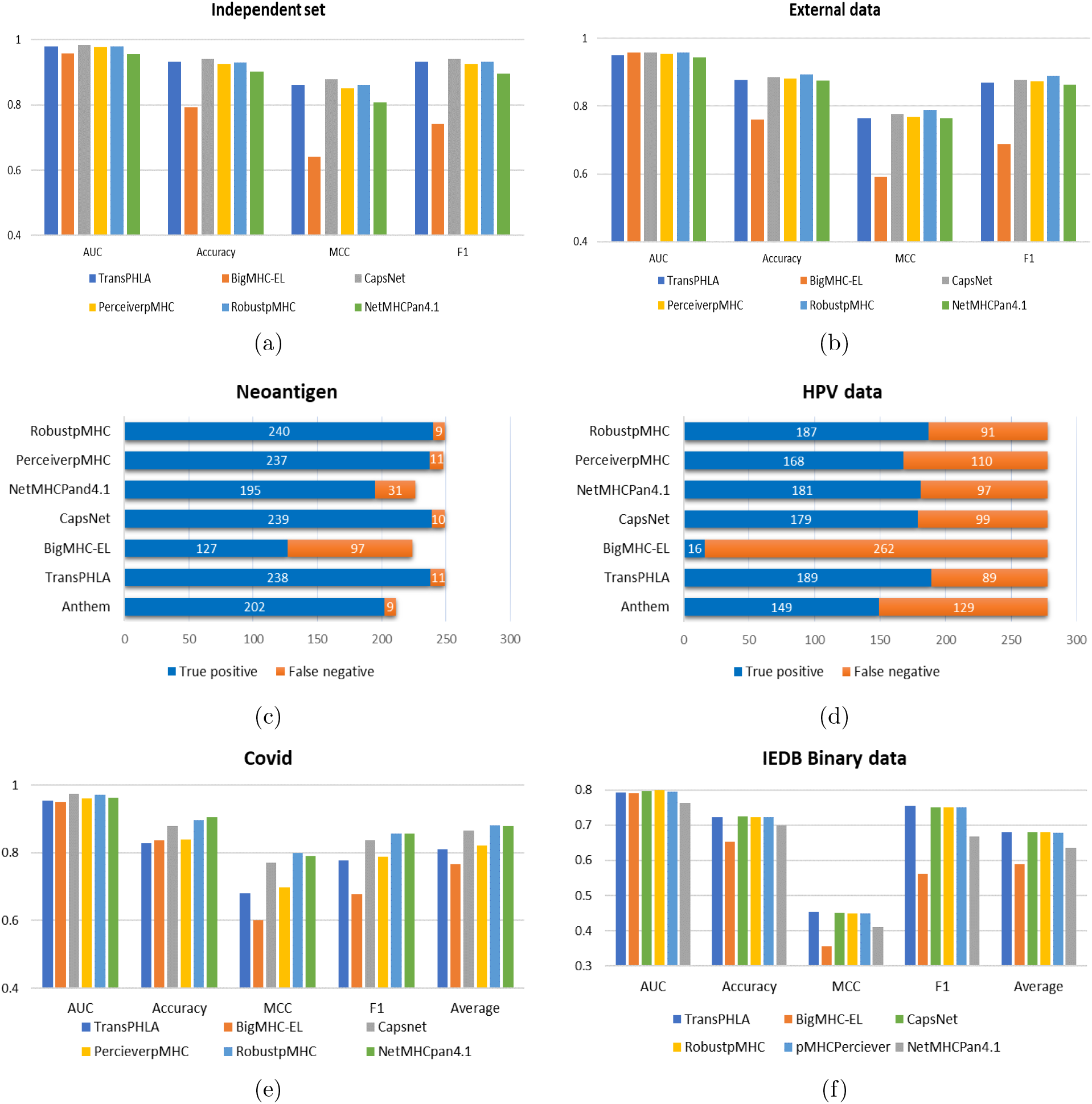
Robustness evaluation. Comparison of RobustpMHC with recent approaches on (a) Anthem Independent set, (b) Anthem External set, (c) Neoantigen data and (d) HPV data, (e) COVID data and (f) Binary set from IEDB database.

### 1.3 Transfer learning evaluation

While the learning based approaches have shown promising results for pHLA binding prediction, their performance steeply deteriorates for prediction on data with significantly different distribution. In Figure 3a-e, we plot the distribution of peptide length across different datasets. It can be seen that distribution of Anthem independent set is very similar to the train set, thus the performance of end-to-end approaches are better as can be seen in Figure 2a. On the other hand, for HPV dataset where the distribution of peptide length is different as can be observed by comparing Figure 3d with Figure 3a, the performance of all approaches significantly deteriorates as depicted in Figure 2d. Hence, for an extensive evaluation of out-of-domain (OoD) generalisation we select two datasets: (i) Neopeptide dataset presented by a very recent method BigMHC [2]. This dataset comprises 198 positive instances and 739 negative instances, and it has served as a benchmark for transfer learning. (ii) We compile a dataset collected from IMGT/3Dstructure-DB dataset [32], which we refer to as CrystalIMGT is an experimental verified Dataset comprising of 14997 crystallographic experimental samples. The peptide length distributions of these datasets are shown in Figure 3e,f respectively. It can be seen from Figure 3f-h that existing approaches suffer from significant performance degradation on these datasets and especially CrystalIMGT.

**Fig. 3.**
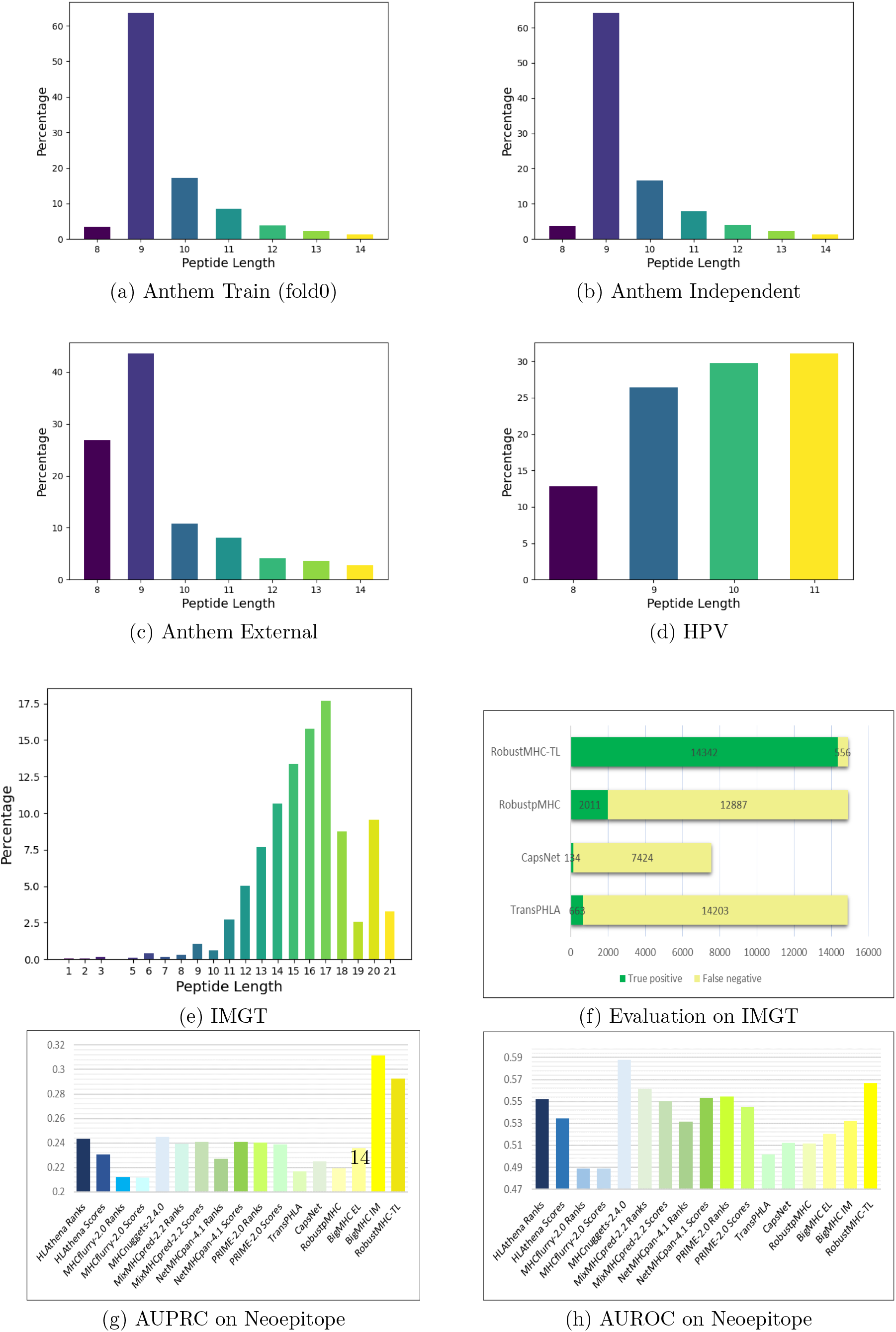
(a-e) Distribution of peptide length for different datasets. It can be seen that when datasets have different distribution like (d) HPV and (e) CrystalIMGT, the performance of existing methods significantly decreases. (f,g,h) Evaluation of transfer learning on (f) CrystalIMGT and (g,h) Neoepi-tope dataset.

To tackle such large distribution shift, we present a transfer learning paradigm for

RobustpMHC. The CrystalIMGT we select 10% of the data randomly and consider them as an independent training set for transfer learning, while for the Neopeptide dataset, we follow the BigMHC and use the independent training and testing sets provided by them for fair evaluation. For transfer learning we keep our feature extraction module i.e. Perceiver IO module (refer Figure4) frozen and retrain the projection module with a smaller learning rate (1*e^−^*^4^). The evaluation results from Figure 3f-h show that this transfer learning paradigm can significantly improve the performance pHLA binding prediction even on novel datasets which are significantly different from the training dataset.

### 1.4 Ablation studies

#### Comparison with data augmentation and contrastive learning

We next evaluate the contribution of robust training, for we compare RobustpMHC without robust training i.e. PerceiverpMHC. Furthermore, it can be argued that whether equation (1) is the most optimal way of robust training? To answer this question we compare our robust training pipeline with two popular self-supervised learning approaches: (i) data augmentation [53] and contrastive learning [36]. In data augmentation the existing dataset is augmented with a synthetic data by transforming the original sample. Here we consider random mutation as a transformation and retrain PerceiverpMHC on the combined dataset, we refer to this model as PerceiverpMHC-Re. (ii) In contrastive learning (specifically noise constrastive learning) for each mini-batch of the training data each sample is passed through a transformation. Then infoNCE loss [36] is used to learn representations robust to a given transformation. Here again considering random mutations as transformation, we train a variant of our model that minimizes infoNCE loss instead of the proposed loss 1. We refer to this variant as PerceiverpMHC-CE. From supplementary Figure S3, it can be seen that our approach (RobustpMHC) consistently outperforms PerceiverpMHC, PerceiverpMHCRe and PerceiverpMHC-CE, which validates the contribution of our loss function in obtaining better generalisation across diverse datasets.

#### Other transformer architectures for full sequence pMHC binding prediction

To validate the selection of Perceiver IO as our base module for encoding peptide-MHC sequence we train the vanilla transformer [54] on full MHC sequence and compare with PerceiverpMHC and another popular efficient transformer architecture Reformer [26] which we call ReformerpMHC. We can observe from the supplementary Figure S2 that PerceiverpMHC performs better than TransformerpMHC (slightly inferior) also being over 4 times faster in both training and evaluation on a single RTX-3090 GPU while the ReformerpMHC performs significantly inferior as can be seen in supplementary Figure S1. It is worth mentioning the for drug discovery a wide range of peptides are evaluated. Moreover before evaluation the peptide sequences are divided into sub-sequences of different lengths of 8-14mers and then each sub-sequence is evaluated against MHC for identifying binding candidates, which makes evaluation speed crucial for practical reasons. This motivates our choice for selecting PerceiverpMHC as our base model.

#### Effect of number of mutations for robust training

Our training pipeline aims to improve the pMHC representation against small amount of mutations in the MHC and peptide sequences. We hypothesize that small amount of mutations on an average does makes the neural network robust, and small mutations can be seen as noise in the data. Whereas, for large mutations the properties of the allele and peptide changes, thus the binding probability should be different. To evaluate this hypothesis we train RobuspMHC on small amount of mutations *m* = *⊓*(0, 0.15) and compare with that trained on large amount of mutations *m* = *⊓*(0, 0.35). It can be observed from supplementary figure S4 that small mutation improves the efficacy of the approach while large mutations degrades the performance.

#### Effect of different peptide length

While several existing approaches are able to work with different peptide lengths their performance noticeably varies as the peptide length changes. This is due to the bias of the training data, since the networks expect similar distributions. Here we explicitly study the effect of peptide length on performance of RobustMHC for Anthem external dataset. it can be seen from supplementary Figure S5, that RobstMHC’s performance drops for peptides with lengths greater than 9, since maximum amount of training data belongs to peptides with lengths 8,9. It is a common issue of leaning based approaches and similar observations have been reported by TranspHLA and CapsNet.

## 2 Method

### 2.1 Datasets

#### Anthem dataset

The Anthem dataset [33] comprises of three types of dataset the training set for model training and model selection, the independent test set and the external test set for model evaluation and methods comparison. The data sources for the training and independent test set are the same (i) four public HLA binders databases ([12], [43], [28] and [41]), (ii) allotype-specific HLA ligands identified by mass spectrometry in previously published studies as described in TransPHLA [10]. The training dataset consists of 112 types of HLA alleles, including 3,59,166 EL and 17,95,830 negative instances. Peptides of negative data are sequence segments that are randomly chosen from the source proteins of IEDB HLA immunopeptidomes. The independent and external set consists of 112 types of HLA alleles with 85,876 positive and 85,562 instances. Because the source of the independent test set and the training set are the same, the data distributions for the training set and independent test set are very similar, TranPHLA [10] used an additional external set consisting of 5 types of HLA alleles with 51,984 positive and 51,881 negative instances was used to perform a fairer evaluation.

#### Neoantigen dataset

The Neoantigen dataset presented by the Chu et al. [10] consists of 250 Neoantigen samples was compiled from experimentally verified non-small-cell lung cancer, melanoma, ovarian cancer and pancreatic cancer pHLA binders.

#### HPV dataset

The human papilloma virus (HPV) dataset presented by Bonsack et al. [8] studies one of the most common sexually transmitted disease. It comprises of 278 experimentally verified pHLA binders from HPV16 proteins E6 and E7, consisting of 8–11-mer peptides. Following [8], we also consider ‘binder’ according to IC50*<*100µM for the HPV vaccine data.

#### COVID dataset

The coronavirus disease 2019 (COVID-19) belongs to family SARS-CoV-2-induced pneumonia, named by World Health Organization as coronavirus disease 2019 (COVID-19), has been declared a pandemic on the 11th of March 2020 since its first appearance in Wuhan, China, in December 2019 [13].

Coronavirus disease 2019 (COVID-19), caused by severe acute respiratory syndrome coronavirus 2 (SARS-CoV-2) infection, became a worldwide pandemic affecting millions of people [47]. Several efforts were made to identify HLA-related susceptibility to SARS-CoV-1 also belong to this family [6] to understand the variation in HLA affect both susceptibility and severity of COVID-19 infection could help identify risk and may support future vaccination strategies. Experimental Dataset collected from IMGT/3Dstructure-DB structure database[32] contains crystallographic experimental data consists of 42 experimentally verified samples.

#### CrystalIMGT dataset

We utilized the IMGT/3Dstructure-DB database [14] to provide a unique resource of expertise with detailed specific annotations on structural data of IG, TR, MHC and RPI, from human and other vertebrate species, extracted from the Protein Data Bank PDB [7]. The IMGT/3Dstructure-DB is publicly available structure database [32] comprise experimentally verified 3D data contains 15395 pHLA binders. IMGT/3Dstructure-DB integrates data from sequence and structural sources and provides identification of IMGT genes and alleles expressed in the IG, TR and MHC with known 3D structures.

#### Neoepitope dataset

For evaluating the transfer learning capability of our approach, we follow the BigMHC [2]. We use the training (positive = 1,407; negative = 4,778), and validation (positive = 173; negative = 515) split for transfer learning our projection block keeping the weights of Perceiver IO block frozen. The transfer learning evaluation dataset consisted of a total of 198 immunogenic and 739 non-immunogenic neoepitopes after removing the intersection with all other pMHC data, were compiled from different sources such as NEPdb [58], Neopepsee [24], TESLA [56], and a data collected from 16 cancer patients using the MANAFEST Albert et al. [2]. They only kept peptides of length at least 8 and at most 11 and peptides with dummy amino acid ‘X’ were removed.

#### MHC-I database

IPD-IMGT/HLA database [4] is the official repository for the WHO Nomenclature Committee for Factors of the HLA System that receives submissions from laboratories in over 46 countries and active website uses from users in over 150 countries. Total 36036 MHC-I sequences across 38 species from the IPD-IMGT/HLA Database version 3.53.0 use for self-supervised learning.

### 2.2 RobustpMHC network architecture

RobustpMHC leverages cross-attention mechanism [54] in latent space to extract features from both peptide and MHC sequences. To ensure computational efficiency, especially with longer sequences, we employ efficient transformer variants, specifically Perceiver IO [17]. Additionally, to train our model to capture features robust to minor mutations, we introduce small random mutations into the peptide and MHC sequences. We then train a student model to be resilient to these perturbations in a self-supervised manner. To predict binding probabilities from these learned representations, we incorporate a non-linear projection block followed by a classifier. The overall architecture of RobustpMHC is depicted in Figure 4, and in the following sections, we will provide a detailed description of each individual building block.

**Fig. 4.**
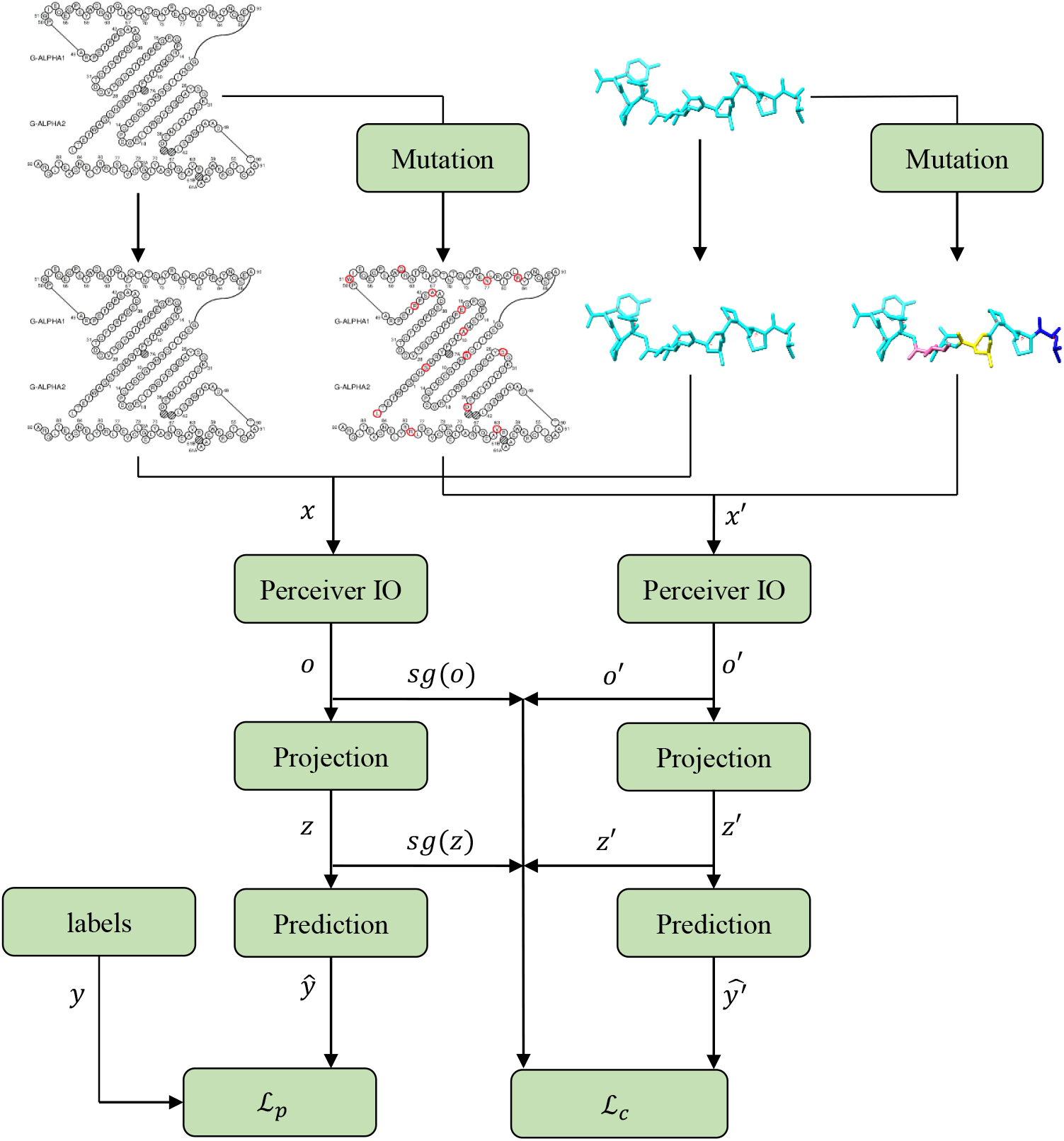
Pipeline of the proposed RobustpMHC approach. Both the MHC and pepdies are mutated with small probability and concatenated for the input sequence. We then train two identical networks containing Perceiver IO blocks for feature extraction, projection and prediction blocks for predicting the binding probability from the Perceiver IO features. We refer to them as teacher (*ϕ*(*·*)) and student network (*ϕ ^′^*(*·*)). The teacher network is receives the original sequence (*x*), while the student gets mutated sequences (*x^′^*). The teacher also has access to true labels (*y*) and can be trained directly by minimising the prediction loss *𝕷p*. We train the student by matching the latent features of the student network [*o ^′^, z ^′^*] with their counterparts of the teacher minimising the consistency loss *𝕷c*. The central idea is to make the student robust to small mutations.

#### Mutation block

The mutation block is inspired from mask language modelling and contrastive self-supervised learning used in natural language processing [23] where randomly an input token is masked, such designs have also shown significant improvement in the robustness in even other domains such as mask-autoencoder [15]. In this work instead, we mutate a token representing embedding of a an amino acid. For each batch we sample a mutation probability between *m* = ⊓(0, 0.15) uniformly for peptide and *m* = ⊓(0, 0.05) uniformly for MHC. Then for each amino acid in the sequence, we independently sample a random variable (*p_i_*) from Bernoulli distribution. We mutate an amino acid (*i*) if *p_i_ < m*. Note that, if we only keep mutation probability (*m*) fixed then this could lead to a distribution shift, since during test time, peptides might not be mutated. Therefore, we also randomly sample the mutation probability (*m*) from a uniform distribution.

#### Perceiver IO block

In this work, we aim to work with entire sequences and showcase that using attention mechanisms deep neural networks can help to learn the representation of full sequence efficiently, as can be seen for Figure 1. To this end, we make our architecture based on efficient transformer architectures such as Perceiver IO [17]. Since, Transformers scale very poorly in both compute and memory they require significantly higher training time for processing full sequences. Perceiver IO uses cross-attention to map large inputs sequences to a smaller number of latent features. Then processing is performed entirely on these latent tokens, and finally they are decoded to an output space as shown in Figure 5(a). Therefore, Perceiver IO has no quadratic dependence on the input or output size: encoder and decoder attention modules depend linearly on the input and output size (respectively), while the latent attention is independent of both input and output [17].

**Fig. 5.**
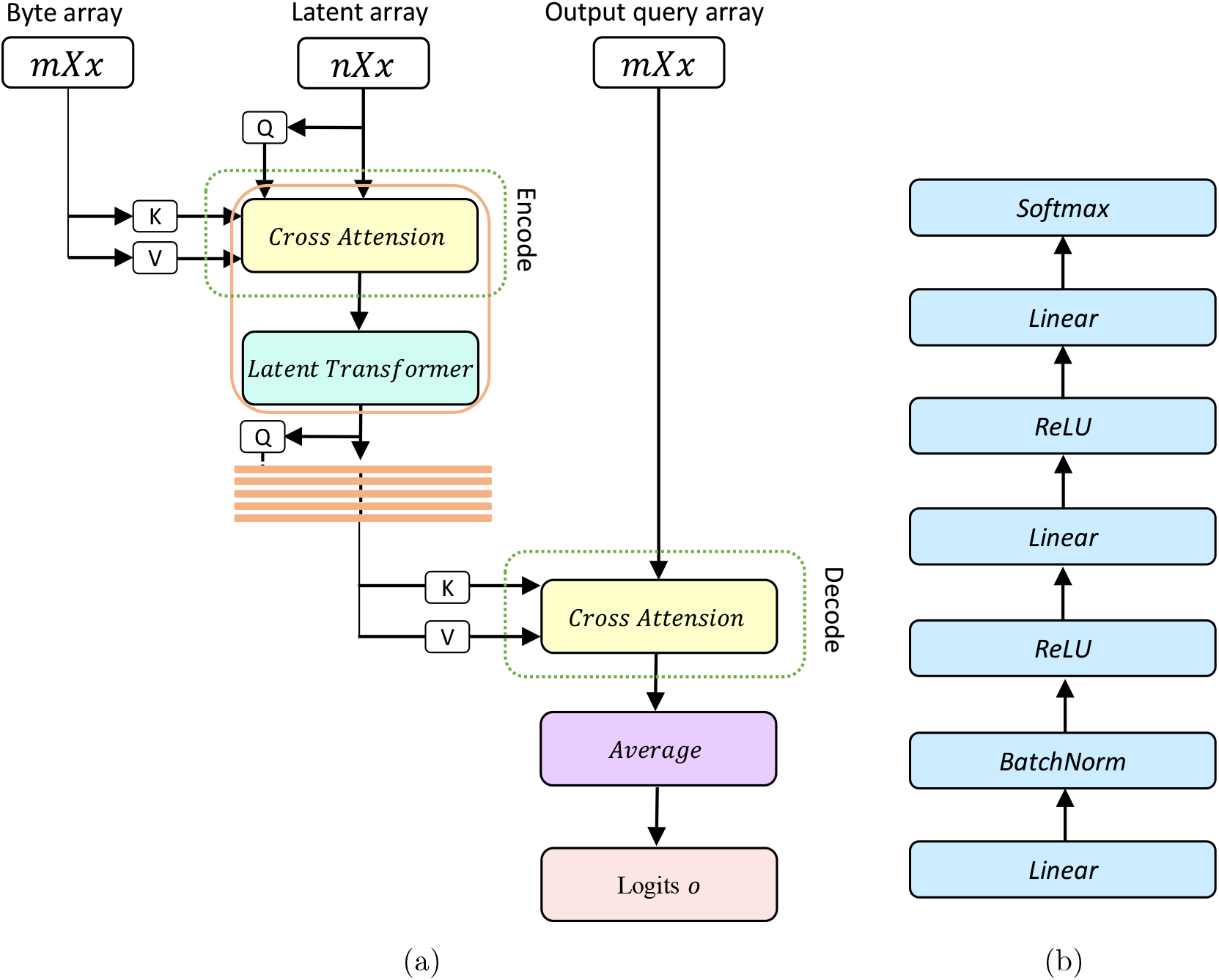
Building blocks of the proposed RobustpMHC approach. (a) Perceiver IO block: It projects the input sequence with large number of tokens to a smaller number of latent tokens using cross-attention and all processing is done in the latent space. (b) Projection block: the projection block in a 3-layer neural network that projects the Perceiver features to classifier space.

#### Projection and Prediction blocks

For obtaining binding scores from the Perceiver IO features, we flatten the latent features from the decoder of the Perceiver IO. These features are then propagated through a 3-layer shallow neural network with dense layers and ReLU activation similar to TransPHLA [10] as shown in Figure 5(b). For prediction we just take the Softmax of the output score of the projection block to obtain the binding probability. Similar to previous approaches [10],[31], we threshold

the binding probability at 0.5 for converting it into class labels.

##### Algorithm 1

**RobustpMHC**

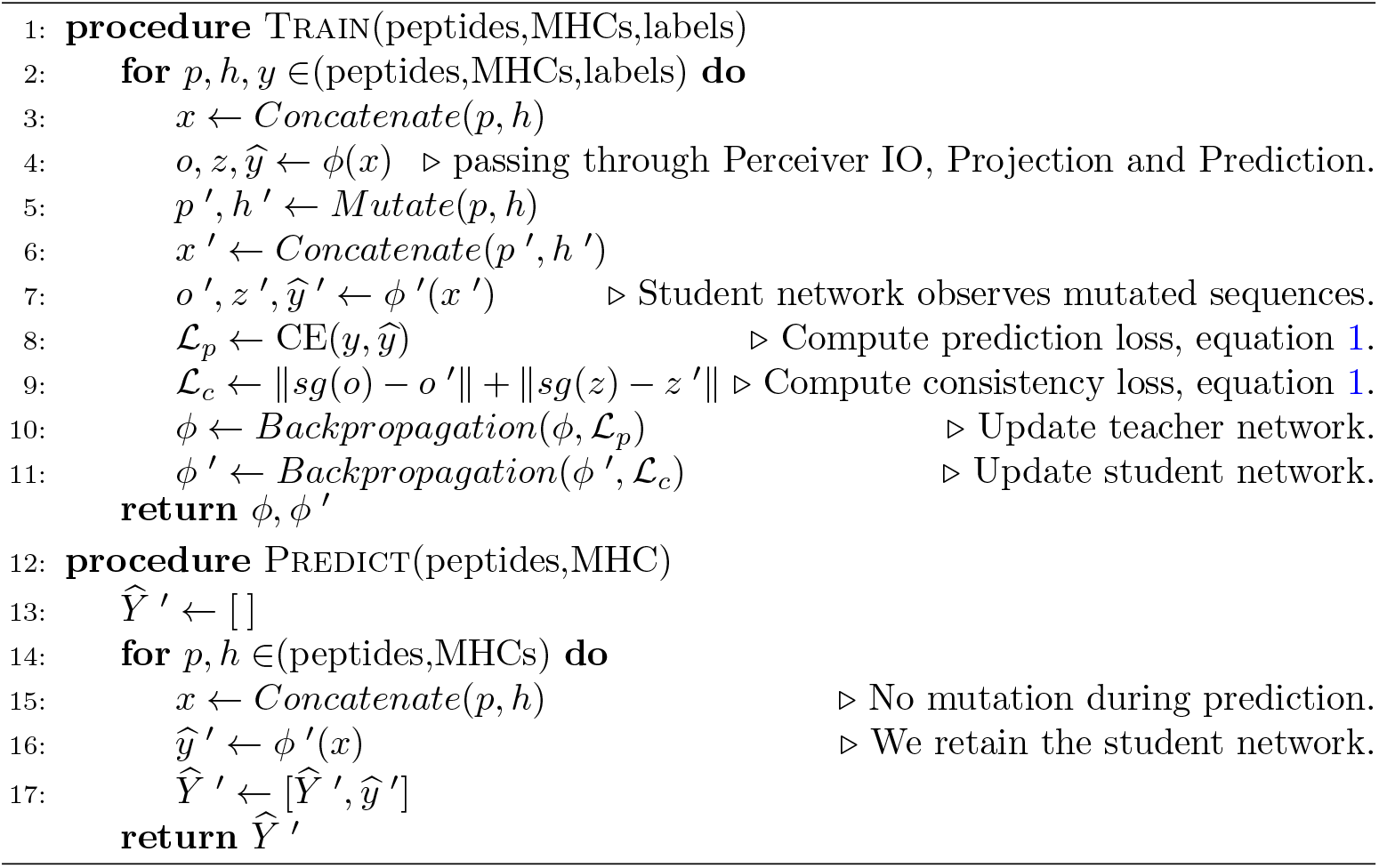

### 2.3 Self-supervision for robust feature learning

In ablation studies section 1.4, we show that simply training the network with mutations or using contrastive loss results in inferior performance. Therefore, we present a self-supervision strategy inspired from knowledge distillation to learn robust features with the help of mutations. Our strategy comprises of training two identical networks together. The first network which we refer to as the teacher (*ϕ*(*x*)) has access to unmasked inputs (*x*) and true class labels (*y*). While the second network, which we call student (*ϕ ^′^*(*x*)) can only observe the mutated input *x ^′^* and has to regress to the latent features [*o, z*] of the teacher network as shown in Figure 4(a). Therefore, our objective function (*𝕷* = *𝕷_p_* + *𝕷_c_*) consists of two losses, the prediction loss (*𝕷_p_*), and the consistency loss (*𝕷_c_*),

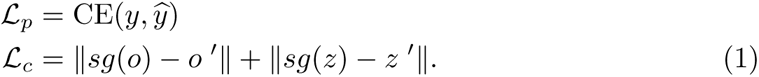

Where, CE is the category-cross entropy loss and *sg*(*·*) is the stop gradient operator that refrains the gradient flow to the teacher during backpropagation so that only student is trained on the gradients of *𝕷_c_*. During the evaluation phase we only consider the predictions from the student networks i.e. 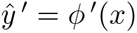. Note that, while evaluating we do not enforce mutation to input sequence. This self-supervision is summarized in algorithm 1.

### 2.4 Implementation Details

We train RobustpMHC on single GeForce RTX 3080 GPU which has 24 GB memory. The model was compiled on PyTorch 1.10.0 and CUDA 11.7 kernel. We train RobustpMHC for 100 epochs and use early stopping retaining the one with best validation performance. Following TransPHLA [10] for fair comparison with previous approaches, we also train five models on the training data where the negative samples of the training set are divided into 5 parts for five-fold cross-validation, however we report the mean performance unlike TranspHLA where best performing model’s performance is reported. We use the batch size of 1024 for training since this is the largest size we could fit on single GeForce RTX 3080 GPU. We employ Adam optimiser for training with learning rate 0.001, we further reduce the learning rate to its one-tenth, when the validation performance is not improved in 4 epochs.

### 2.5 Compared methods

We exhaustively evaluate our approach by conducting two types of case studies, on one hand we compare our approach on six different datasets for validating robustness and on the other hand, we evaluate transfer learning capability of our approach on two datasets. During these studies we compared with the state-of-the-art methods such as TranspHLA, CapsNet, the IEDB recommended method (NetMHCpan-4.1), over fifteen IEDB baseline methods and transfer learning based method (BigMHC) published very recently, RobustMHC achieves most consistent performance across all experiments. We here explain few selected recent approaches and highlight their contrast with RobustMHC. It is worth noting that aprat from our approach only DeepAttentionPan can take full sequences, other approaches require pseudo sequences.

**NetMHCpan4.1** [44] is a popular tool for simultaneously predicting BA and EL. This method consists of an ensemble of 100 single-layer networks, each of which consumes a peptide 9-mer binding core and a subsequence of the MHC molecule. The 9-mer core is extracted by the model, whereas the MHC representation, termed a ‘pseudo-sequence,’ is a predetermined 34-mer core extracted from the MHC molecule sequence. The 34-mer residues were selected based on the estimated proximity to bound peptides so that only residues within 4 ^°^A were included.

**MHCflurry-2.0** [37] is an ensemble of neural networks that predicts BA and EL for MHC-I. BA prediction is the output of a neural network ensemble, where each member is a two- or three-layer feed-forward neural network. Then, an antigen processing convolutional network is trained on a subset of the BA predictions, along with the regions flanking the N-terminus and C-terminus, to capture antigen processing information that is missed by the BA predictor. EL prediction is the result of logistically regressing BA and antigen processing outputs.

**TransPHLA** [10] is a transformer-based model that adopts NetMHCpan pseudo-sequences for MHC encoding. This model encodes peptides and MHC pseudo-sequences using the original transformer encoding procedure before inferring the encodings via rectified linear unit (ReLU)-activated fully connected layers. TransPHLA was trained on the Anthem data for binary binding prediction.

**MHCnuggets** [49] comprises many allele-specific LSTM networks to handle arbitrary-length peptides for MHC-I and MHC-II. Transfer learning was used across the alleles to address data scarcity. MHCnuggets trained on qualitative BA, quantitative BA, and EL data. MHCnuggets trained on up to two orders of magnitude fewer data than the other methods.

**HLAthena** [48] uses three single-layer neural networks trained on mass spectrometry data to predict presentation on short peptides with length in the range [8,11]. Each of the three networks trained on a separate peptide encoding: one-hot, BLOSUM62 [3], and PMBEC40. In addition, the networks consumed peptide-level characteristics, and also amino acid physiochemical properties. The outputs of these networks were used to train logistic regression models that also accounted for proteasomal cleavage, gene expression and presentation bias. HLAthena also saw performance gains when considering RNA-seq as a proxy for peptide abundance.

**CapsNet** [21] proposes a capsule neural networks architecture for learning the representation of Anthem’s datasets. CapsNet-MHC architecture, is built upon four major units: a) data encoder, b) feature extractor, c) binding dependencies extractor, and d) binding predictor. They used CNNs with the attention mechanisms for feature extraction from the encoded peptide and MHC sequences, using Blosum62 [3] input encoding method.

**BigMHC** [2] present transfer learning method on the prediction of immunogenic neoepitopes. Initially, the model is pre-trained on publicly available data of MHC class I peptide presentation. Subsequently, the model undergoes a transfer learning process, where it leverages this pre-existing knowledge to fine-tune its predictions for immunogenic neoepitopes.

**DeepAttentionpan** [19] propose an improved pan-specific model, based on convolutional neural networks and attention mechanisms for more flexible, stable and interpretable MHC-I binding prediction.

NetMHCpan, HLAthena and MHCflurry rely on ensemble of shallow neural networks for binding prediction. The ensemble help in reducing variance however shallow neural networks are not able capture the fine nuances in pMHC sequence, which are crucial for binding prediction. MHCnuggets on the other hand considers pMHC pseudo-sequences and employs a LSTM to extract the latent features however, LSTM features face issues in capturing longer sequence dependencies. Therefore, TransPHLA employs a single layer Transformer and CapsNet utilises a Capsulenet architecture for extracting better features. While these approaches rely hand crafted psuedo-sequences, we show that deep attention networks are able to capture similar features from full sequences. DeepAtentionPan incorporated attention layers between convolution layers for extracting features from full sequences, however their performance is significantly inferior to recent pseudo-sequence methods. We use efficient tranformer models based on latent cross-attention to efficiently extract better features, we further propose self-supervision to make the models generalise better across diverse datasets. BigMHC presents a transfer learning paradigm to predict binding on immunology datsets, we further show that the features learned by RobustpMHC also transfers well to immunology datatset (we showcase better performance as compared to BigMHC on one of the two metrics). Moreover, on other datasets BigMHC’s performance is far inferior as compared to ours. Hence, our approach can tackle full sequences and is able to show consistently better results across diverse datsets.

## 3 Discussion

### Conclusion

In this research, we have demonstrated the efficacy of IMGT/RobustpMHC model in predicting peptide-HLA class I binding probabilities particularly when considering full MHC sequences. IMGT/RobustpMHC belongs to the category of generalized pan-specific models that are not restricted by HLA alleles or peptide length. Our model employs cross-attention mechanisms within deep neural networks, therefore exhibits capability to learn comprehensive representations of MHC sequences, underscoring the potential of efficient transformer architectures like Perceiver IO in computational immunology. Furthermore, we also incorporated self-supervised learning by training the network with mutations that empowered our model to capture subtle inter-dependencies between peptide and HLA sequences, enabling it to generalize effectively and resulting in significant improvements in binding probability predictions. Notably, our model maintains robust performance even in the presence of mutations, which is vital for real-world applications. Collectively, the combination of these techniques propels our Perceiver IO-based model to the forefront of peptide-HLA binding prediction, offering a promising tool for immunotherapy and vaccine design. We believe that our model highlights the importance of harnessing advanced deep learning techniques in tackling complex biological problems.

### Limitations

While IMGT/RobustpMHC consistently demonstrates improved performance compared to the state-of-the-art approaches across a wide range of datasets, it still relies on additional transfer learning data for CrystalIMGT and Neopeptide datasets. Notably, the performance of all approaches remains suboptimal on these specific datasets. Consequently, we are convinced that the field of peptide-MHC (pMHC) prediction could benefit from extensive pre-training and domain adaptation techniques to bridge this performance gap, and IMGT/RobustpMHC represents a crucial step in this direction.

Furthermore, a common issue we encounter across all datasets is the generation of negative samples for training these approaches, which is typically done randomly without experimental verification. Consequently, these negative samples may contain false negatives. We acknowledge that this is one of the shortcomings of existing approaches, including our own, and we intend to address this issue in our future research endeavors.

### Webserver availability

The webserver is available at https://www.imgt.org/RobustpMHC/

### Code availability

Once this paper is accepted, the implementation code and trained models will be available at (https://github.com/anjanakushwaha/RobustpM HC). PyTorch 1.10.0 and CUDA 11.7 was used to calculate performance metrics. Pandas v.2.1.1 and Numpy v.1.22.3 were used for data processing. Matplotlib v.3.5.1 were used to generate figures.

### Data availability

Once this paper is accepted, the data will be available at (https://github.com/anjanakushwaha/RobustpMH

C/tree/main/dataset). It contains the training data, independent test data, external test data, anthem dataset, neoantigen dataset, HPV dataset, COVID dataset, crystalIMGT dataset and neoepitope dataset.

## Supplementary information

The online version contains supplementary material available.

## Supporting information

Ablation studies

## Acknowledgments

We express our gratitude to the entire IMGT® team for their ongoing dedication and unwavering enthusiasm. IMGT® is currently supported by the Centre National de la Recherche Scientifique (CNRS) and the University of Montpellier. IMGT® is member of the French Infrastructure ”Institut Fraņcais de Bioinformatique”, IFB as well as member of BioCampus, MAbImprove and IBiSA. AK was co-funded by “Direction de la Recherche, du Transfert Technologique et de l’Enseignement Supérieur” of Occitanie Region under the N° 20007399 / ALDOCT-001023 contract as well as IMGT® proper resources. This work was granted access to the High Performance Computing (HPC) resources of Meso@LR and of “Centre Informatique National de l’Enseignement Supérieur” (CINES).

## Notes

### Competing Interest Statement

The authors have declared no competing interest.

