## Supplementary material for "IMGT/RobustpMHC: Robust Training for class-I MHC Peptide Binding Prediction": Ablation studies

### 1. EVALUATION MATRICES

In this section, we describe the metrics used for evaluating our approach and comparison with other recent approaches in the paper for completeness.

### A. F1

The F1 score is a commonly used metric in machine learning, particularly in binary classification tasks. It provides a balance between precision and recall, two important evaluation metrics.
**Precision** is the ratio of true positives (TP) to the sum of true positives and false positives (TP + FP). It measures the accuracy of positive predictions made by the model. A high precision indicates that the model makes fewer false positive errors.

$$\text{Precision} = \frac{TP}{TP + FP}$$

**Recall** is the ratio of true positives to the sum of true positives and false negatives (TP + FN). It measures the ability of the model to correctly identify all positive instances in the dataset. A high recall indicates that the model makes fewer false negative errors.

$$\text{Recall} = \frac{TP}{TP + FN}$$

The F1 score is the harmonic mean of precision and recall, and it combines both metrics into a single value. An F1 score of 1 indicates perfect precision and recall and 0 indicates poor performance in terms of both precision and recall.

$$\text{F1 Score} = \frac{2}{\frac{1}{Precision} + \frac{1}{Recall}}$$

Overall, the F1 score provides a balanced evaluation of a model's performance in binary classification tasks, making it a valuable metric for various applications.

#### B. ROC (The Receiver Operator Characteristic)

An ROC curve, or receiver operating characteristic curve, is like a graph that shows how well a classification model performs. It helps us see how the model makes decisions at different levels of certainty. The curve has two lines: one for how often the model correctly identifies positive cases (true positives) and another for how often it mistakenly identifies negative cases as positive (false positives). By looking at this graph, we can understand how good the model is and choose the threshold that gives us the right balance between correct and incorrect predictions. The Receiver Operator Characteristic (ROC) curve is an evaluation metric for binary classification problems. It is a probability curve that plots the TPR against FPR at various threshold values and essentially separates the 'signal' from the 'noise.' In other words, it shows the performance of a classification model at all classification thresholds.
This curve plots two parameters:
**True Positive Rate (TPR)** is a synonym for recall:

$$\text{TPR} = \frac{TP}{TP + FN}$$

**False Positive Rate (TPR)** is :

$$\text{FPR} = \frac{FP}{FP + TN}$$

#### C. AUC(Area Under the ROC Curve)

It measures the overall performance of the binary classification model. As both TPR and FPR range between 0 to 1, So, the area will always lie between 0 and 1, and A greater value of AUC denotes better model performance.The AUC is calculated by computing the area under the ROC curve.

#### D. Accuracy

Accuracy simply measures how often the classifier correctly predicts. We can define accuracy as the ratio of the number of correct predictions and the total number of predictions.Accuracy provides an intuitive and easily interpretable measure of a model's performance. However, it may not always be the best metric to use, especially in cases of imbalanced datasets where one class significantly outnumbers the other. In such cases, a model that predicts the majority class most of the time can have a high accuracy, but it may not be a useful or reliable classifier.

$$\text{Accuracy} = \frac{TP + TN}{TP + TN + FP + FN}$$

#### E. MCC (The Matthews Correlation Coefficient)

The Matthews Correlation Coefficient (MCC) is a metric used for evaluating the performance of binary classification models, particularly in situations where the classes may be imbalanced. It takes into account true positives (TP), true negatives (TN), false positives (FP), and false negatives (FN) to provide a balanced assessment of the model's performance.The MCC ranges from -1 to +1, where +1 indicates a perfect classification, 0 indicates random classification, and -1 indicates complete disagreement between predictions and actual labels.

$$\text{MCC} = \frac{TP \times TN + FP \times FN}{\sqrt{(TP + FP)(TP + FN)(TN + FP)(TN + FN)}}$$

#### F. AUPRC (The area under the precision-recall curve)

The area under the precision-recall curve (AUPRC) is a useful performance metric for imbalanced data in a problem setting where you care a lot about finding the positive examples. If your model achieves a perfect AUPRC, it means your model found all of the positive examples (perfect recall) without accidentally marking any negative examples as positive (perfect precision). The "average precision" is one particular method for calculating the AUPRC. It provides a comprehensive assessment of a model's ability to distinguish between the positive and negative classes.

### 2. ABLATION STUDY RESULTS

Here we present the results for Ablation studies conducted in the paper answering the research questions regarding the choice of efficient transformer architecture, loss functions, mutation probability and effect of peptide length.

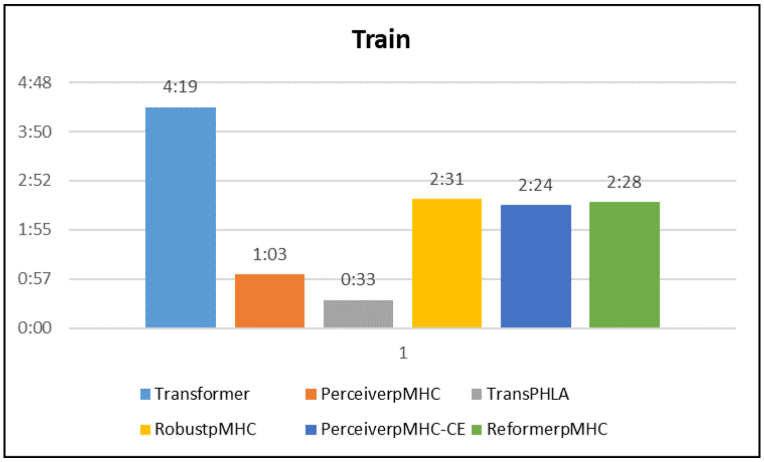


(a)

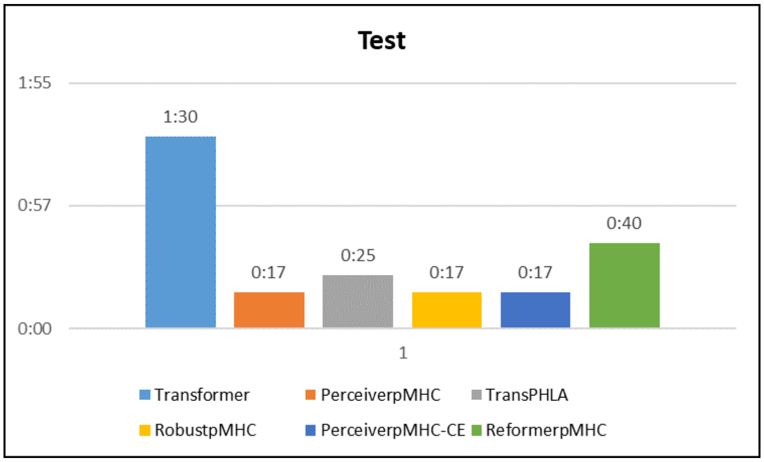


(b)

**Fig. S1. Timing comparison.** Training and Testing time in minutes and seconds comparing RobustpMHC with other transformer variants on (a) Anthem training dataset fold-0 for 1 epoch. (b) Anthem External dataset fold-0 for 1 epoch. It can be seen that PercieverpMHC is over four time faster than Transformer making it suitable candidate for processing full sequences.

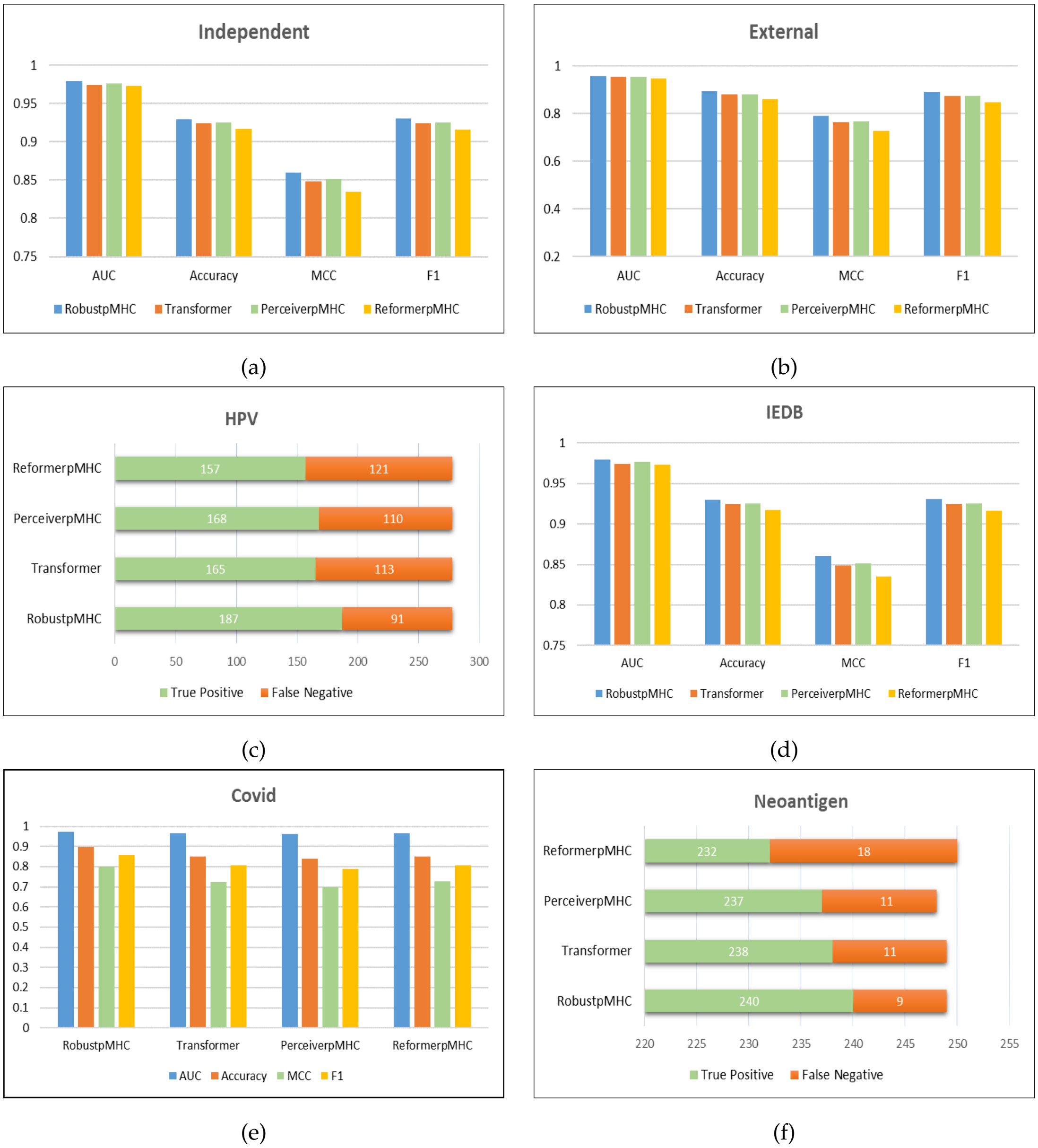


**Fig. S2. Alternative transformer architectures study.** Comparison of RobustpMHC with different transformer architectures such as Transformer, ReformerpMHC, PerceiverpMHC across six datasets. It can be seen that PercieverpMHC is at par (slightly better) with Transformer while being much faster.

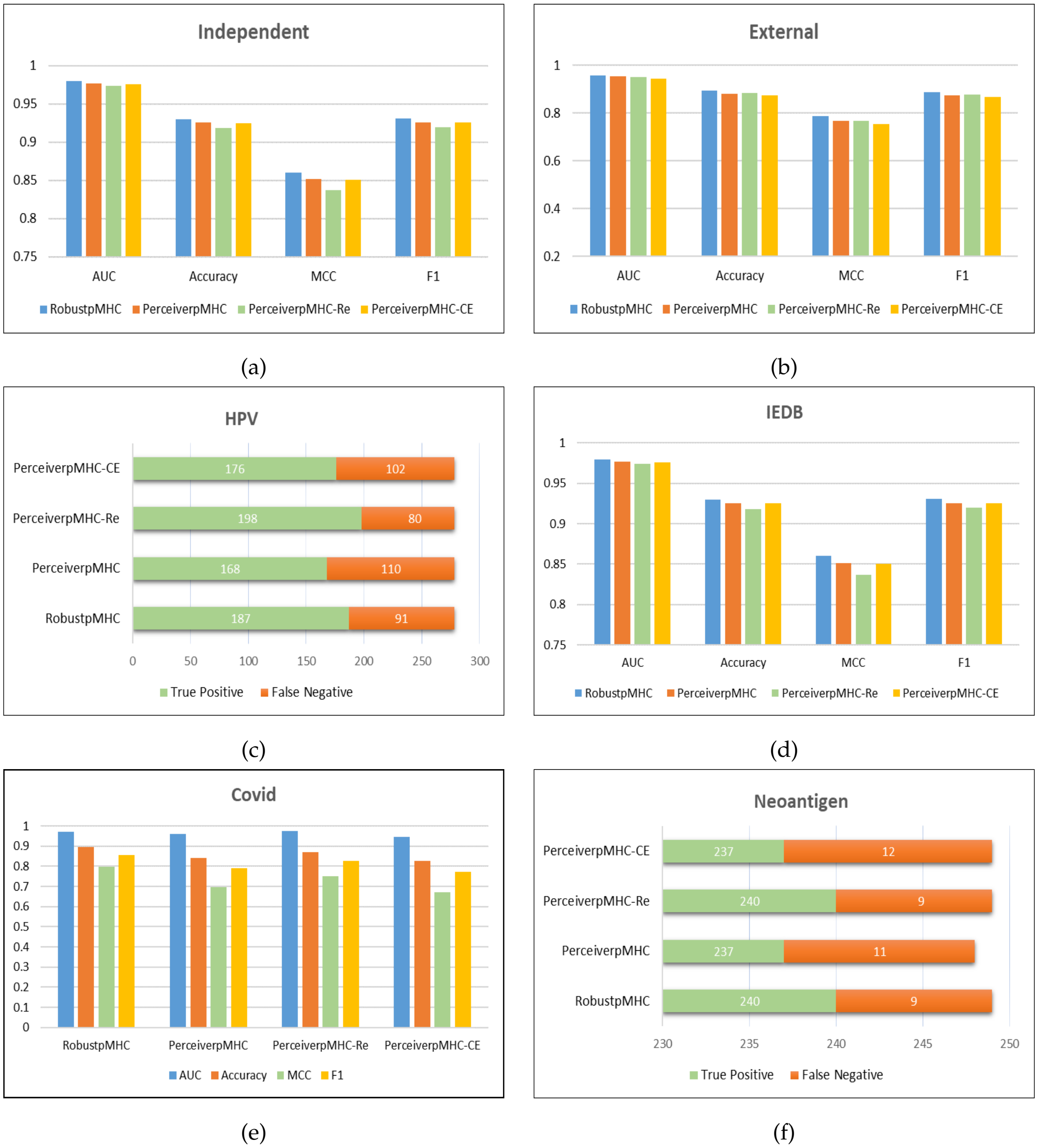


**Fig. S3. Self-supervised learning methods study.** Comparison of RobustpMHC's training approach with no self-supervision (PerceiverpMHC), data augmentation (PerceiverRe) and contrastive learning (PerceiverpMHC-CE) approaches across six datasets. Here we use mutations as data augmentation. It can be seen that RobustpMHC outperforms all other baselines, which justifies the choice of our loss function.

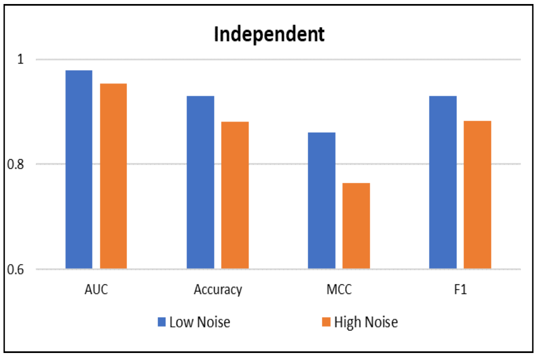


(a)

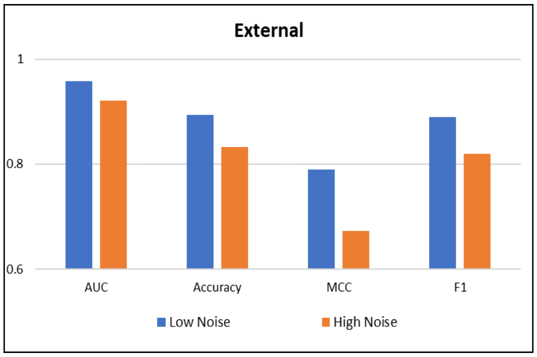


(b)

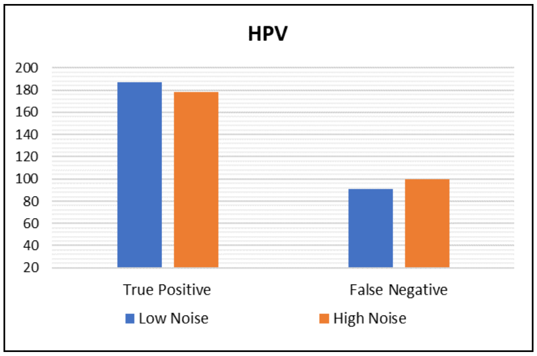


(c)

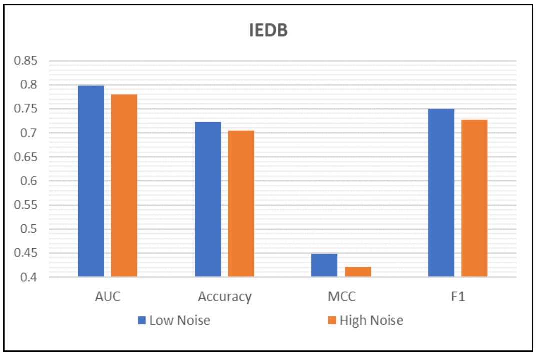


(d)

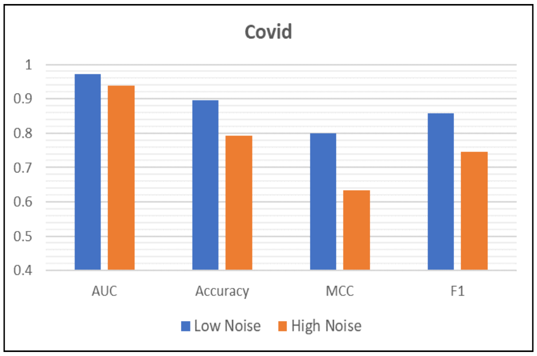


(e)

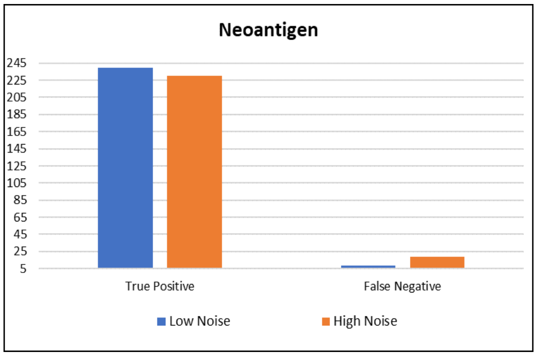


(f)

**Fig. S4. Mutation amount evaluation.** Comparing the small mutation based robust training with large mutation based robust training on 6 datasets. While small amount of mutations help in achieving better generalization capability. It can be seen that, if we make the mutation probability large then performance deteriorates because for large mutation at an average the properties changes, therefore the binding probability might change.

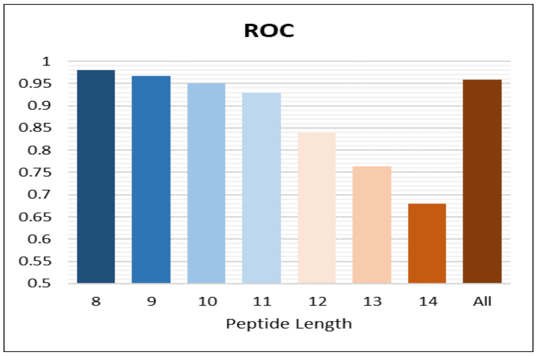


(a)

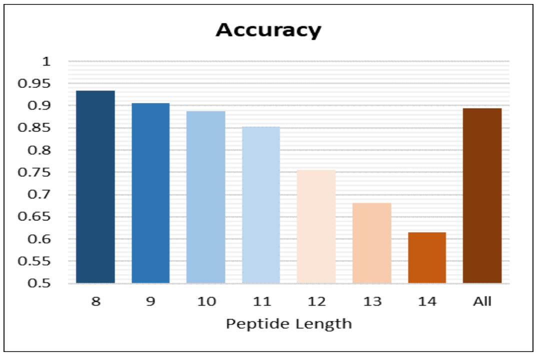


(b)

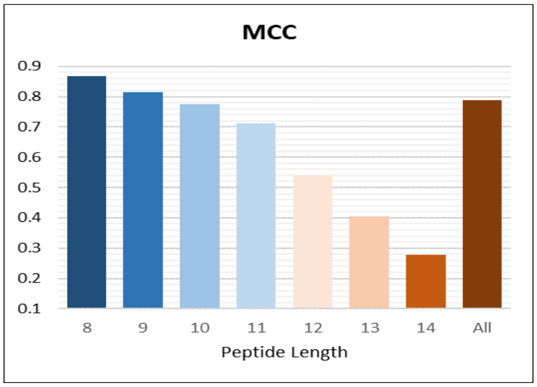


(c)

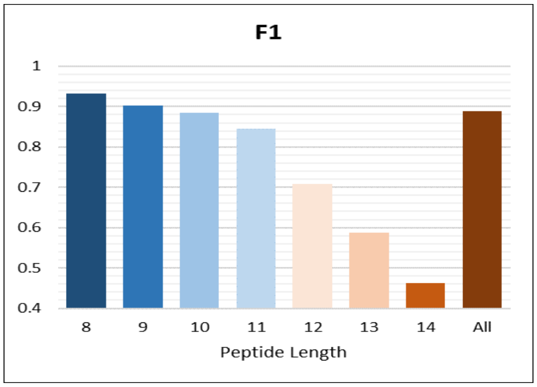


(d)

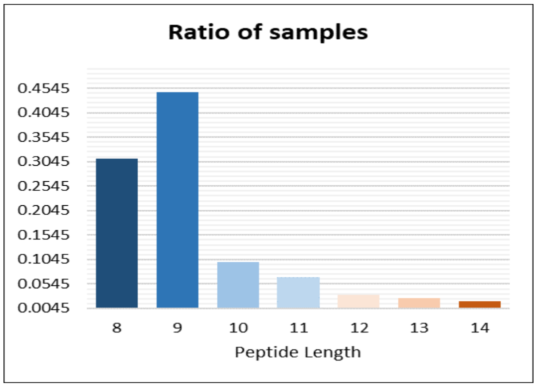


(e)

**Fig. S5. Peptide Length evaluation.** Performance of RobustpMHC on different peptide lengths for four metrics (a) ROC, (b) Accuracy (c) MCC (d) F1 on Anthem external dataset. (e) Distribution (Probability mass function) of external dataset based on peptide lengths. The last bar in (a-d) represents the overall performance i.e. on the entire dataset including all peptides. Notice the correlation between Data distribution and performance as the length of peptide varies.
